# Human and bovine serum albumin, but not mouse serum and egg-white albumin, promote reactivation of viable but non-culturable *Mycobacterium tuberculosis* via the activation of protein kinase-dependent cell division processes

**DOI:** 10.1101/2021.11.22.468319

**Authors:** Yuta Morishige, Yoshiro Murase, Kinuyo Chikamatsu, Hiroyuki Yamada, Akio Aono, Yuriko Igarashi, Keisuke Kamada, Yoshiko Shimomura, Makiko Hosoya, Satoshi Mitarai

## Abstract

We investigated the mechanisms underlying the fetal bovine serum (FBS)-induced reactivation of the viable but non-culturable (VBNC) state of *Mycobacterium tuberculosis* (Mtb) using the H37Rv strain. We found that bovine serum albumin (BSA), a major component of FBS, could reactivate the VBNC state of Mtb cells without other reactivation-promoting agents. Human serum albumin also had a restorative effect similar to BSA, but mouse serum albumin and egg-white albumin (ovalbumin) did not, suggesting that a common protein structure between bovine and human serum albumin is essential for reactivation. In addition, antioxidative agents, such as N*-*acetyl-*L*-cysteine, showed no restorative effect, suggesting that the restoration of culturability might not be due to the antioxidative property of BSA. BSA-mediated reactivation is inhibited by H89 and staurosporine. These inhibitors are known to inhibit multiple protein kinases, including serine/threonine protein kinases, in Mtb, suggesting the involvement of mycobacterial protein kinases in cell division during reactivation. These findings provide new insights into the mechanisms of Mtb reactivation, and may contribute to the development of novel strategies for the treatment of latent tuberculosis.

**Importance:** Reactivation of dormant, including viable but non-culturable (VBNC) *Mycobacterium tuberculosis* (Mtb) cells, represents a critical step in the progression from latent infection to active tuberculosis. However, the molecular mechanisms that trigger this transition are not yet fully understood. We herein report that bovine serum albumin (BSA), a major component of fetal bovine serum, can restore the culturability of VBNC Mtb cells. Human serum albumin showed a similar effect, whereas mouse and egg-white albumins did not, suggesting that the specific structural features shared between bovine and human albumins are essential for reactivation. The lack of reactivation by the antioxidant N-acetyl-*L*-cysteine indicates that the mechanism is not related to redox balance. Inhibition of BSA-induced reactivation by H89 and staurosporine implicates mycobacterial serine/threonine protein kinases in this process. These findings highlight the previously unrecognized role of albumin-kinase signaling in Mtb reactivation and suggest new molecular targets for preventing tuberculosis relapse.

## Introduction

*Mycobacterium tuberculosis* (Mtb) develops a latent phenotype, dormancy, due to various stresses, such as host immune response, oxygen depletion, and nutritional starvation, *etc.* The Dormant Mtb cells can persist inside the host and may establish latent tuberculosis infection (LTBI), which requires activation. Lowering the risk of Mtb activation in patients with LTBI is an important strategy to reduce the incidence of TB (1). The viable but nonculturable (VBNC) state is a variant of dormancy in non-sporulating bacteria (2). VBNC cells are a non-culturable population that can be induced by exposure to certain stressors. They cannot grow on nutrient media even after the removal of stress and require a much longer time or specific stimulus for growth (2, 3). It is increasingly being recognized that the VBNC state may play an important role in the survival of Mtb in human hosts.

Many pathogenic bacteria, including Mtb, can enter the VBNC state and are considered to be in a protective state in the host environment (4). Various *in vitro* models, such as models of hypoxia, nutritional starvation, a lipid-rich environment, and other multiple stresses, have been reported to induce a VBNC state in Mtb cells (5–9). A deeper understanding of the activation mechanisms of these cells is necessary for the prevention of active tuberculosis.

One of the major hypotheses underlying the formation of VBNC bacteria involves the oxidative damage generated by harsh external conditions (10). Some studies have reported that antioxidation during culture is one of the key mechanisms by which VBNC bacteria are reactivated (11, 12). Some mycobacterial culture media containing bovine serum albumin (BSA) and BSA are considered to be involved in the detoxification of growth-suppressing agents that are spontaneously generated during culture (13–15). However, few studies have focused on the effect of such media on the reactivation of VBNC cells or as an antioxidative agent in mycobacterial cultures.

Previously, the NADH oxidase inhibitor diphenyleneiodonium chloride (DPI) was reported to simply and rapidly induce a VBNC state in Mtb H37Ra, and incubation with fetal bovine serum (FBS) facilitated the reactivation of VBNC cells, whereas incubation with oleic-albumin-dextrose-catalase (OADC) supplementation did not (16). This report provides important clues for constructing a simple and rapid assay system to reactivate VBNC Mtb cells. However, we found that the whole body of FBS might not be essential for the reactivation of DPI-induced H37Rv VBNC cells, but the presence of BSA, a major component of FBS, seemed to be sufficient to reactivate the VBNC state, as observed for both H37Rv and H37Ra. Because FBS contains a high concentration of BSA, we hypothesized that BSA would act as a reactivation-promoting agent for DPI-induced VBNC cells.

One possible reactivation mechanism of BSA is its antioxidative properties, which have been well studied (17). Another possibility is cellular processes involving cAMP, an important second messenger in the cell. Shleeva *et al*. showed that the presence of high amounts of cAMP provided by adenylyl cyclase is essential for the reactivation of *M. smegmatis* and Mtb from the VBNC state (18, 19). Notably, overexpression of Rv*2212* gene, which encodes adenylyl cyclase in Mtb (20), significantly affects both entry into and reactivation from the VBNC state (19).

Understanding the detailed reactivation mechanism of VBNC is crucial for reducing the risk of Mtb disease development. In this study, we reported that the reactivation-promoting mechanisms of BSA against DPI-induced VBNC Mtb might not be due to its antioxidative property or the activation of adenylyl cyclase.

## Results

### I. DPI treatment induced a VBNC state in Mtb H37Rv

Yeware *et al.* previously reported that DPI treatment, which inhibits the electron transport system, resulting in the inhibition of aerobic respiration, could effectively induce the VBNC state in Mtb H37Ra cells (16). In our study, we confirmed that H37Ra and H37Rv cells could effectively enter the VBNC state with DPI treatment, as evaluated by culturability, loss of acid-fastness, and accumulation of neutral lipids. Figure 1 shows the DPI treatment results for the induction of the VBNC state in H37Rv cells. As shown in Fig. 1(A), the numbers of culturable cells in the untreated control population and DPI-treated population were significantly different: 4.43 ± 1.43 × 10^8^ CFU/mL for the untreated control population and 4.64 ± 6.10 × 10^3^ CFU/mL for the DPI-treated population (*p* = 0.033). Fig. 1(B) shows that the proportion of CFDA-positive cells in the untreated control and DPI-treated populations was not significantly different (81.8 ± 11.4% vs 65.9 ± 11.8%, *p =* 0.169). Representative images of CFDA/EtBr dual-stained cells from the untreated control and DPI-treated populations are shown in Fig. 1(C). These results indicate that the majority of DPI-treated H37Rv cells were alive, while their culturability deteriorated significantly.

**Figure 1.**
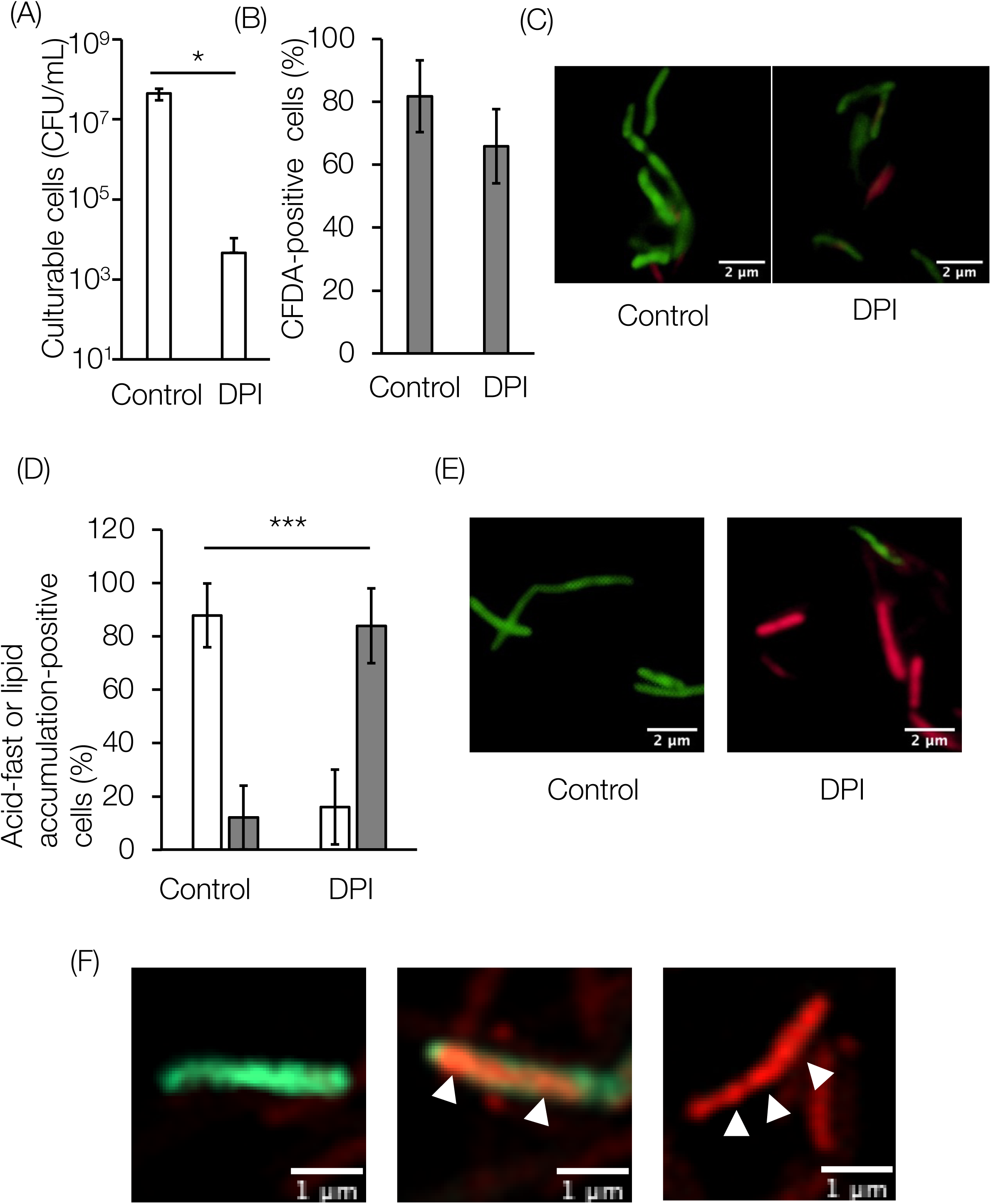
Viability analyses of DPI-treated Mtb H37Rv. (A) Number of culturable cells after DPI treatment. The CFU/mL values were determined by plating the cells onto 7H10 plates in duplicate. Asterisks indicate statistically significant differences (\**p* < 0.05, Welch’s *t*-test). Data represent mean ± SD from three independent experiments. (B) Percentage of esterase-active cells after DPI treatment. Approximately 850 cells were counted directly using a 100× objective lens. Data represent mean ± SD from three independent experiments. (C) Fluorescence micrographs of Mtb H37Rv cells stained with carboxyfluorescein diacetate (CFDA) and ethidium bromide (EtBr). Cells stained with CFDA (green) were esterase-positive, and those stained with EtBr (red) were membrane-damaged. Representative images from three individual experiments are shown. Scale bar: 2 µm (D) Percentage of acid-fast-positive cells (open column) or acid-fast-negative but lipid accumulation-positive cells (gray column). Approximately 1,100 cells were counted directly using a 100× objective lens. Data represent mean ± SD from four independent experiments. Asterisks indicate significant differences (\*\*\**p* < 0.005, Welch’s *t*-test). (E) Merged fluorescence micrographs of Mtb H37Rv cells stained with auramine-O and Nile Red. Cells stained with auramine-O (green) were acid-fast-positive; those stained with Nile Red (red) were acid-fast-negative but lipid accumulation-positive. Representative images from three individual experiments are shown. Scale bar: 2µm (F) Magnified views of Airyscan confocal super-resolution microscopy observations of three different Mtb cells obtained from DPI-treated cultures. Only acid-fast-positive cells (left), both acid-fast-positive and lipid accumulation-positive cells (center, counted as acid-fast-positive), and acid-fast-negative cells with only lipid accumulation-positive cells (right). Dual-positive cells probably represent a transition to a non-culturable state. The indicated foci with arrowheads represent the neutral lipid bodies. Scale bar: 1 µm

Auramine-O/Nile Red dual staining showed a reduction in acid fastness and accumulation of neutral lipids. As shown in Fig. 1(D), the proportion of acid-fast-negative/lipid accumulation-positive cells was 84.0 ± 14.0% in the DPI-treated population, and 12.1 ± 11.9% in the untreated control population (*p* < 0.01). Representative images of auramine-O/Nile Red dual-stained cells in the untreated control and DPI-treated populations also showed distinctive differences in the proportions of acid-fast-positive cells and lipid-accumulation-positive cells (Fig. 1[E]). The loss of acid-fastness was also confirmed by Ziehl-Neelsen staining; the majority of DPI-treated populations were found to have lost their acid-fastness (Suppl. Fig. 1). Interestingly, Airyscan microscopy revealed that some DPI-treated cells were positive for acid-fast and neutral lipid staining, suggesting the presence of cells in the transitional state from culturable to VBNC (Fig. 1[F]). These results indicate that DPI treatment induced the VBNC state in H37Rv and H37Ra.

### II. BSA is essential on the reactivation of DPI-induced VBNC Mtb cells

To evaluate whether DPI-treated H37Rv cells could regain culturability by supplementing the medium with FBS, we performed a reactivation assay on DPI-treated H37Rv cells supplemented with FBS or OADC. As shown in Fig. 2(A) and (B), our results partially support the previous study (16) with some differences; not only were the DPI-treated H37Rv cells reactivated by incubation with FBS-supplemented medium, but also the OADC-supplemented medium. Addition of albumin alone to the medium can also induce reactivation. The addition of FBS to the BSA-free Dubos medium slightly reduced the regrowth rate; however, cells were successfully reactivated at the end of incubation with or without FBS. This phenomenon was also observed for H37Ra (Suppl. Fig. 2). Thus, we used H37Rv for further analyses in this study.

**Figure 2.**
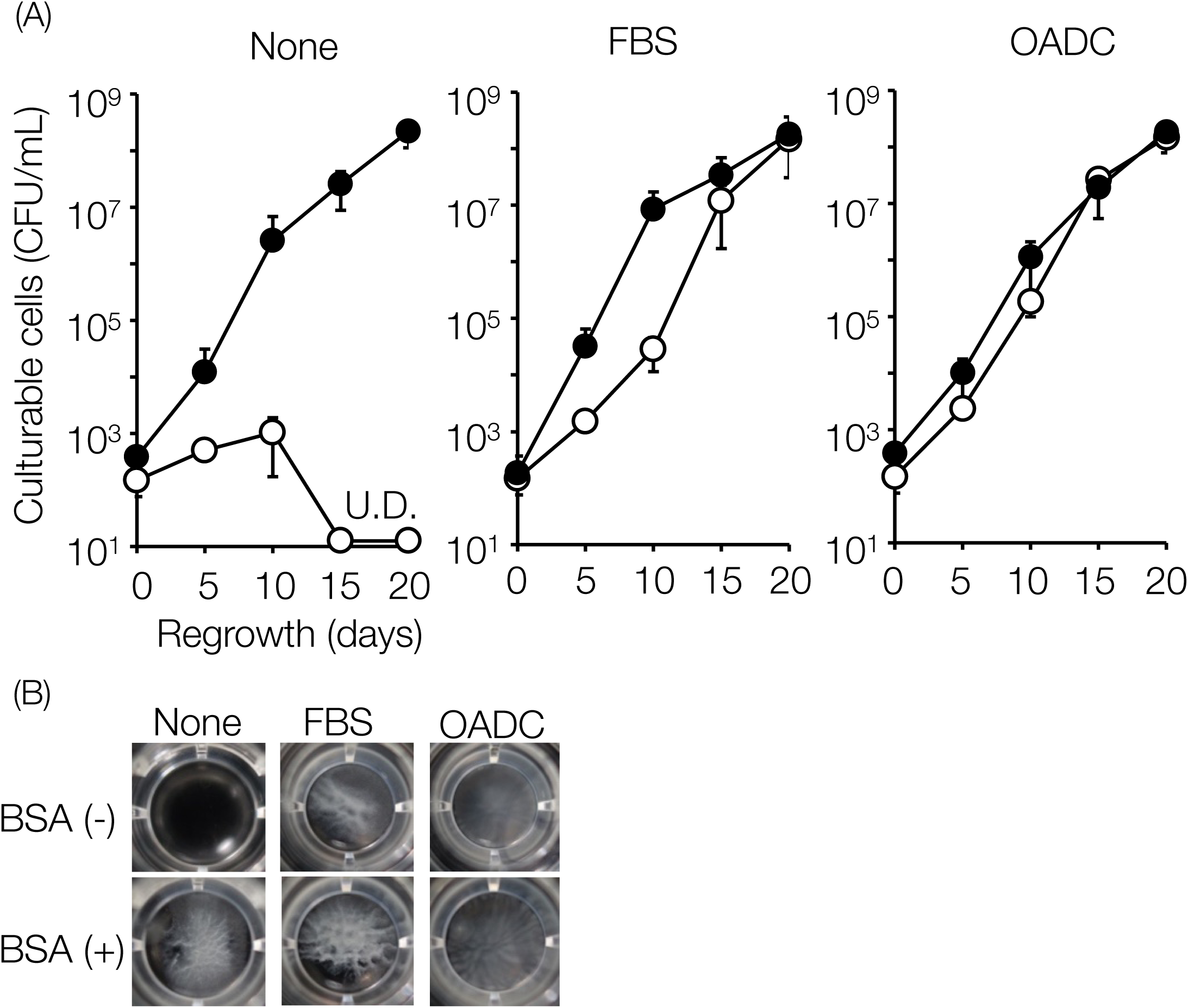
Reactivation of DPI-treated Mtb H37Rv cells by FBS and OADC supplementation. (A) Number of culturable cells on the indicated days of incubation, without supplementation, or supplemented with 2% FBS or 10% OADC in the presence (closed circle) or absence (open circle) of 0.1% BSA. The CFU/mL values were determined by plating the cells onto 7H10 plates in duplicate. Data represent mean ± SD from three independent experiments. U.D. indicates the limit of detection (20 CFU/mL). (B) Representative images of regrowth under the indicated conditions were captured at the end of incubation. None, FBS, and OADC represent no supplementation, 2% FBS supplementation, and 10% OADC supplementation, respectively.

Incubation with sodium pyruvate, which has been reported to have a reactivation-promoting effect on VBNC cells elsewhere (21–23), led to transient regrowth by day 15 and a slight reduction at day 20. (Suppl. Fig. 3[A] and [B]).

We also confirmed that the growth rates under the two conditions did not differ significantly (*p* = 0.428 on day 5, Suppl. Fig. 4[A]). Supplementary Figure 4(B) also shows sufficient cell growth. The cells were more aggregative without BSA; however, aggregation was easily dispersed by pipetting, and there was no significant difference in the OD600 value. Albumin is not essential for Mtb growth.

### III. Reactivation-promoting effect by BSA is not attributed to its antioxidative property

It is well known that antioxidants are effective as reactivation-promoting agents for bacteria in the VBNC state (3). We hypothesized that the reactivation-promoting effect of BSA might be due to its antioxidative properties and assumed that human serum albumin (HSA), which has a similar structure, and ovalbumin (OVA), which has a different structure but a stronger antioxidative effect, as well as NAC (antioxidant) and D-mannitol (free radical scavenger), would also reactivate VBNC Mtb. As shown in Fig. 3(A) and (B), the reactivation-promoting effect of albumin was specific to BSA and HSA. Contrary to our expectation, OVA did not show a reactivation-promoting effect but maintained the number of culturable cells in this system. Contrary to our expectations, neither NAC nor D-mannitol showed reactivation-promoting activity.

**Figure 3.**
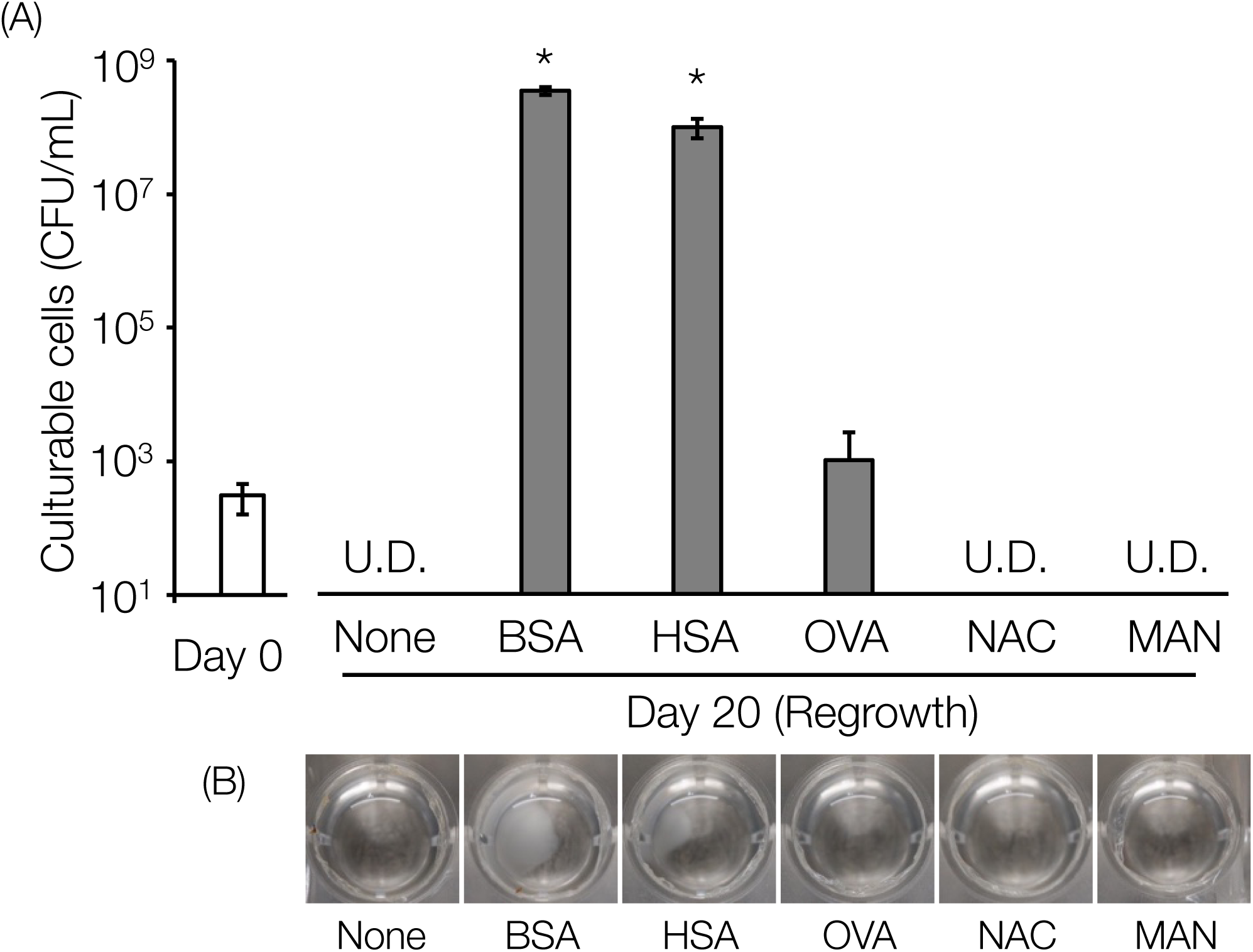
Effect of human serum albumin, ovalbumin and antioxidants on the reactivation of DPI-treated Mtb. (A) The culturable cells at the end of incubation were supplemented with albumins or antioxidative agents. The CFU/mL values were determined by plating the cells onto 7H10 plates in duplicate. Data represent mean ± SD from three independent experiments. Asterisks indicate statistically significant differences between days 0 and 20 by Student’s *t-*test. (\**p*<0.05) (B) Representative images of regrowth captured at the end of incubation. Abbreviations: BSA, bovine serum albumin; HSA, human serum albumin; OVA, ovalbumin; NAC, N-acetyl-L-cysteine; MAN, D-mannitol. The final concentrations of each reagent were as follows: BSA, HSA, and OVA, 0.1% (w/v); NAC, 500 µM; and MAN, 50 mM.

Interestingly, we found that mouse serum albumin (MSA) did not show a reactivation promoting effect (Suppl. Fig. 5). Although only a subset of commercially available species was analyzed, these results suggest that the reactivation promoting effect of serum albumin on DPI-treated Mtb may be specific to certain mammalian species.

We also checked whether the purity of albumin affected the promotion of reactivation using fatty acid- and globulin-free BSA and confirmed that there was no significant difference (*p =* 0.986) in the reactivation-promoting effects (Suppl. Fig. 6). These results suggest that commercially available albumin, including fatty acids, does not affect reactivation of DPI-treated Mtb.

### IV. The reactivation-promoting effect of albumin was canceled by treatment with eukaryotic protein kinase inhibitor

As Shleeva *et al.* reported that cAMP production from adenylyl cyclase is essential for the reactivation of VBNC Mtb (18), we hypothesized that the inhibition of adenylyl cyclase and its downstream enzyme, protein kinase, could modulate the reactivation-promoting effect of BSA. As shown in Fig. 4(A) and (B), SQ22536, an adenylyl cyclase inhibitor, only suppressed reactivation at a very high concentration (10 mM). In contrast, H89, a eukaryotic protein kinase inhibitor, suppressed reactivation in a dose-dependent manner. In detail, incubation with 10 µM or 30 µM H89 resulted in a CFU/mL value for 3.42 × 10^6^ ± 7.45 × 10^5^ CFU/mL and 3.63 ± 1.70 × 10^2^ CFU/mL, while incubation without H89 resulted in 3.00 × 10^8^ ± 5.07 × 10^7^ CFU/mL. We also confirmed that staurosporine, a known mycobacterial protein kinase PknB inhibitor (24), suppressed reactivation at a concentration of 10 µM and resulted in a CFU/mL value of 2.09 × 10^3^ ± 7.62 × 10^2^ CFU/mL, whereas incubation without staurosporine resulted in 3.58 × 10^8^ ± 5.78 × 10^7^ CFU/mL (Fig. 4[A] and [B]). None of the inhibitors used in this study reduced the growth of intact Mtb cells (Suppl. Fig. 7A and 7B).

**Figure 4.**
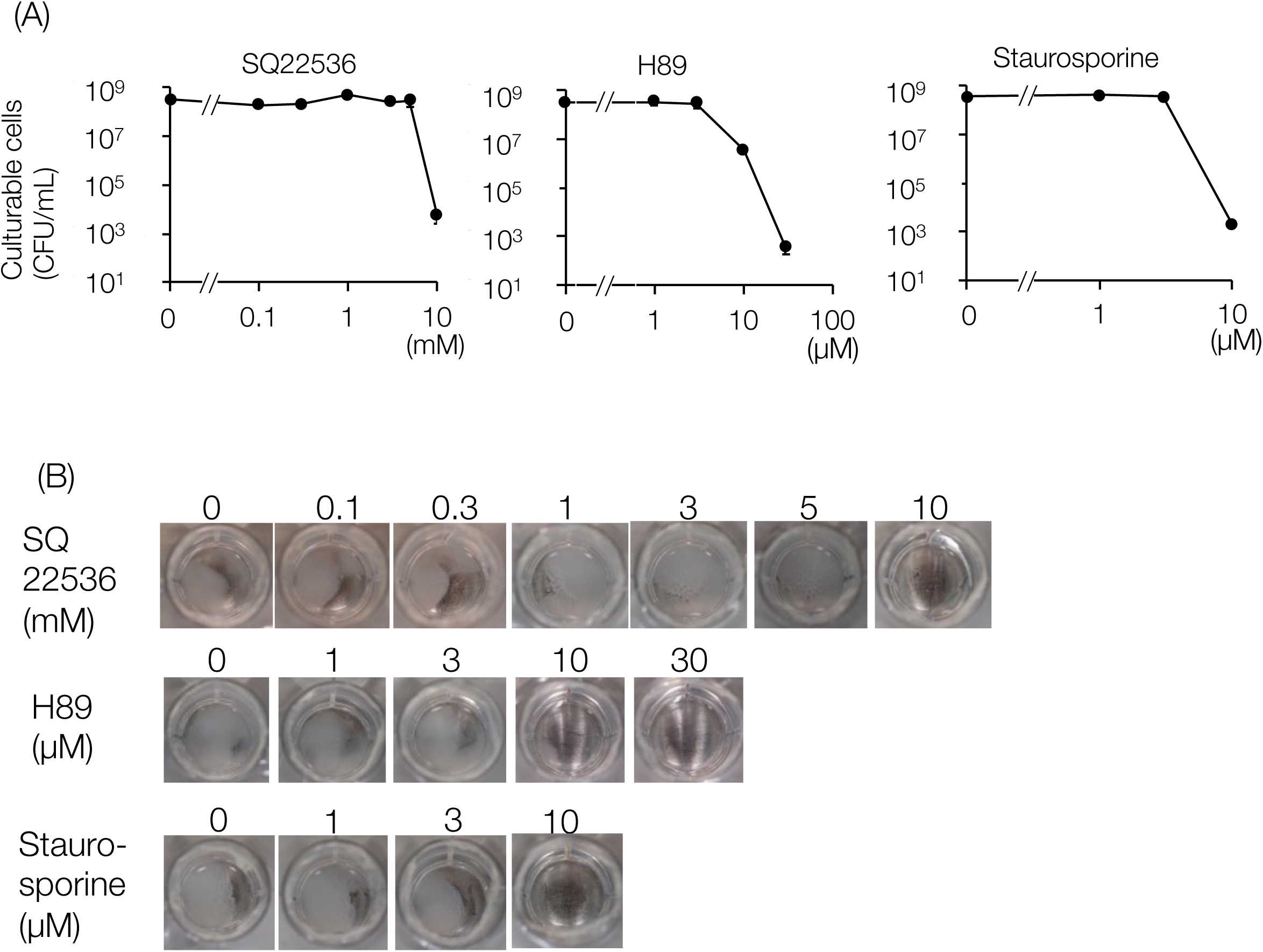
The effect of SQ22536 or H89 or staurosporine on the reactivation of DPI-treated Mtb. (A) Number of culturable cells at the end of incubation, supplemented with 0.1% BSA, SQ22536, H89, or staurosporine. Inhibitors were added at the indicated concentrations. (B) Representative images of regrowth captured at the end of incubation. The CFU/mL values were determined by plating the cells onto 7H10 plates in duplicate. Data represent mean ± SD from three independent experiments.

Since both SQ22536 and H89 are targeted to eukaryotic enzymes, we also performed a molecular docking simulation of these inhibitors against their equivalent targets on Mtb to validate the inhibition assay. As shown in Supplemantary Table 1, SQ22536 and H89 have sufficient affinity for Rv2212 (adenylyl cyclase) and PknA (protein kinase), respectively.

## Discussion

In this study, we found that BSA, a supplement to mycobacterial culture, could act as a reactivation-promoting agent of the VBNC state of Mtb. We also showed that this effect of serum albumin might be species-specific: bovine and human serum albumin had the effect, while mouse serum albumin did not. The predictive mechanism of BSA-induced reactivation was also proposed to involve the activation of mycobacterial protein kinase, not its antioxidative property or the activation of adenylyl cyclase.

The mechanism underlying the effect of DPI is thought to involve the inhibitory effect of NADH oxidase, which results in inhibition of the electron transport system of Mtb. This may induce an unbalanced NADH/NAD ratio and hypoxic response in liquid culture, resulting in a significant decrease in culturability (Fig. 1[A]) (16). However, viability was highly retained (65.9%), suggesting that the majority of the cells were transformed into VBNC (Fig. 1[B]). VBNC transformation was confirmed by auramine-O and Nile Red staining. The loss of acid-fastness and accumulation of neutral lipids are distinctive features of VBNC mycobacterial cells.

Our data also demonstrated that the majority of DPI-treated Mtb H37Rv cells were stained with Nile Red alone (Fig. 1[D]). Airyscan super-resolution microscopy showed the presence of cells stained with auramine-O, which contained an intracellular lipid body stained with Nile Red (Fig. 1[F]). This could also reveal the distribution of the lipid body in the cell as some foci of relatively strong signals of Nile Red, suggesting a transformation from a growing state to VBNC, with drastic alterations in lipid metabolism.

We also found that DPI-induced VBNC could facilitate reactivation not only by incubation with FBS, but also with OADC supplementation and BSA alone, suggesting that albumin might act as a reactivation-promoting agent in both H37Rv and H37Ra (Fig. 2 and Suppl. Fig. 2). These findings were contrary to those of a previous study, which showed that only FBS can facilitate reactivation (16). We considered that the reason for this difference may be due to a contaminant in Tween 80: the presence of an excessive amount of oleic acid. We also confirmed that reactivation was not facilitated by the use of prediluted Tween 80 for the preparation of Dubos medium (unpublished data) for both H37Rv and H37Ra, probably due to extended hydrolyzation of Tween 80 (25). Thus, our results suggest that the key reactivation-promoting agent in FBS is serum albumin, and further analyses were performed using H37Rv. It should be noted that not all cells completely transitioned to the VBNC state in this study.

In general, *in vitro* cultured cells are thought to be a non-uniform population due to the difference in microenvironment; oxygen concentration and nutritional conditions might vary at the local level, which could make the variable subpopulations in metabolic state, as Yeware *et al.* also indicated (16). In our study, a small subpopulation (10^3^ CFU/mL) was found that could not proliferate on Dubos broth without BSA but could proliferate on agar medium (Fig. 2). The results of acid-fast staining also suggested the presence of cells that appeared to be in the process of transitioning from a culturable state to a VBNC state (Fig. 1F). These results suggest that, although the VBNC state could be induced by DPI treatment, the level of induction might vary from cell to cell. Therefore, the results of an *in vitro* VBNC model should be carefully interpreted and the final conclusions must be verified in *in vivo* model with albumin administration.

Our results showed that the reactivation-promoting effect of albumin might be structure-specific because OVA did not facilitate reactivation, whereas BSA and HSA did. Regarding their amino acid sequences, BSA and HSA share 76% sequence homology (26); however, BSA and OVA share less sequence homology (16%). One of the considerable roles of albumin is as an antioxidative agent because of its cysteine residues with free thiols (17, 27). BSA and HSA have only one cysteine residue with a free thiol, whereas OVA has four cysteines with free thiols (28). Although OVA therefore seems to be more reductive than BSA and HSA, their reactivation-promoting effect is paradoxical, indicating that antioxidative properties might not be essential for this effect. This hypothesis was considered to support the reduced effect of NAC and D-mannitol. Moreover, this paradoxical result confirmed that certain common structures of BSA and HSA may be essential. Additionally, we would like to note that MSA did not show the reactivation promoting effect, suggesting that this effect might have species-dependency (Suppl. Fig. 5). At this point, we have found that there are certain differences between bovine and mouse serum albumins (internal data) and analyzed their contribution to this property for the subsequent study, including measurement of the reactivation-promoting effect of MSA on VBNC *M. microti*, a pathogen of small field rodents (29). These results suggest that the effect of albumin may be due to its complex functions rather than antioxidative properties.

Serum albumin is known to act as a good carrier for many biologically active substances, including free fatty acids. It is also known as a detoxifier of growth-arresting substances in culture medium. BSA is added to the mycobacterial culture medium as both a carrier and detoxifier of growth-supporting and growth-arresting substances, respectively. Shleeva *et al*. showed that several types of unsaturated free fatty acids, including oleic, linoleic, and arachidonic acids, can promote the reactivation of VBNC *M. smegmatis* in modified Sauton’s medium, and oleic acid also works in Mtb (18). It should be noted that albumin impurities did not affect reactivation in this study. Although the underlying mechanism is still unclear, both fatty acid- and globulin-free BSA and BSA Cohn fraction V showed a similar reactivation-promoting effect toward DPI-treated Mtb (Suppl. Fig. 6).

The reactivation inhibition assay using SQ22536, H89, and staurosporine provided an important clue for understanding the effect of BSA. In the present study, we suppressed the reactivation of DPI-induced VBNC cells by inhibiting adenylyl cyclase at a very high concentration of SQ22536. Although the underlying mechanism is unclear, adenylyl cyclase may not play a key role in reactivation from a DPI-induced VBNC state, whereas the inhibition of protein kinase by H89 and staurosporine clearly suppressed reactivation. These results suggest that protein kinase might be involved in the reactivation of VBNC Mtb cells triggered by BSA, and that the contribution of adenylyl cyclase may be smaller than that of protein kinase. Mtb encodes eukaryotic Ser/Thr protein kinases including PknA, PknB, and PknD-L. Among them, PknA regulates major cell processes, regulates several types of proteins involved with mycobacterial cell division and morphogenesis via phosphorylation of such proteins in coordination with PknB (30, 31); phosphorylation of GlmU (Rv*1018c*) and MurD (Rv*2155c*), Wag31 (Rv*2145c*), HupB (Rv*2986c*) and ParB (Rv*3917c*), influence the production and cross-linking of peptidoglycan precursors (32–35), the spatial localization of peptidoglycan synthesis(36), DNA condensation and partitioning (37, 38), respectively. Notably, the phosphorylation of FtsZ and FipA by PknA controls cell division under oxidative stress(39, 40). In addition, mycobacterial protein kinase has cyclic AMP-dependent protein kinase domains, which are very similar to the eukaryotic version, which was shown in PknB (41), and staurosporine, which acts as a PknB inhibitor, could also inhibit the reactivation of DPI-induced VBNC Mtb (Fig. 4).

Our study suggests that the inhibition of mycobacterial protein kinase by H89 and staurosporine critically affects several important cellular processes that underlie reactivation (Suppl. Fig. 8). To confirm the role of mycobacterial protein kinases and related proteins in the reactivation of DPI-induced VBNC cells by albumin, further studies involving gene knockdown using CRISPR interference (CRISPRi) (42) are necessary. This is particularly relevant because most of these proteins, except FipA (Rv*0019c*), are essential for *in vitro* growth; (43–46) therefore, genetic manipulation using conventional methods is not permissible.

## Conclusion

Our data showed that the presence of BSA in Dubos medium not only depleted growth-suppressing substances, but also triggered the reactivation of the DPI-induced VBNC state in both H37Rv and H37Ra strains. The reactivation-promoting effects of other kinds of albumins and antioxidative agents, including NAC and D-mannitol, were tested; however, only bovine and human serum albumin showed a reactivation-promoting effect. The results of the inhibition assay of reactivation using SQ22536, H89, and staurosporine suggest that the inhibition of mycobacterial protein kinase suppressed reactivation to a much greater extent than adenylyl cyclase, indicating the presence of a reactivation-promoting protein kinase-dependent pathway. To the best of our knowledge, few studies have focused on the reactivation-promoting effects of albumin on VBNC Mtb. Taken together, our findings indicated that the reactivation of Mtb from a DPI-induced VBNC state could be obtained by interaction with bovine serum albumin and the mechanism of albumin-induced reactivation may involve the activation of mycobacterial protein kinase, which regulates several important processes of cellular component construction and cell division. Although it remains unclear how albumin interacts with VBNC cells, our study provides some important clues for understanding the physiology of the mycobacterial VBNC state.

## Author Contributions

YMori, YMura, and SM conceived and designed the experiments. Y. Mori performed the experiments. YMura, KC, AA, YI, YS, MH, KK, HY, AT, and SM assisted with the experiments. YMori acquired the data, and YMori and YMura analyzed the data. YMori performed a molecular docking simulation and statistical analysis. YMori, YMura, and SM wrote the manuscript. All authors have contributed to the manuscript and approved the submitted version.

## Supporting information

suppl_revised

## Data availability

All data contained in this study can be obtained from the corresponding author upon reasonable request.

## Acknowledgments

The authors thank Dr. K. Suenaga and Mr. H. Sakuma (Carl Zeiss Co., Ltd., Tokyo, Japan) for their assistance in obtaining super-resolution images. The authors also thank Dr. A. Osugi (Research Institute of Tuberculosis, Japan Anti-Tuberculosis Association) for the valuable discussions. The authors also thank Dr. Brian Quinn (Japan Medical Communication). Inc.) for their help with the preparation of this manuscript.

## Materials and Methods

### Bacterial strains, growth media and culture conditions

This study was carried out using *Mycobacterium tuberculosis* H37Rv (ATCC 27294) and H37Ra (ATCC 25177). Both cells were cultured for 3–5 days in a 60 mL glycol-modified polyethylene terephthalate (PETG) bottle containing 20 mL Dubos broth (2.5 g/L Na2HPO4, 2.0 g/L L-asparagine, 1.0 g/L KH2PO4, 0.5 g/L pancreatic digest of casein, 0.2 g/L Tween 80, 0.5 mg/mL CaCl2•2H2O, 0.1 mg/mL CuSO4, 0.1 mg/mL ZnSO4•7H2O, 0.05 g/L ferric ammonium citrate, 0.01 g MgSO4•7H2O) with 5% (v/v) glycerol, and 10% (v/v) ADC supplementation (5.0 g bovine serum albumin fraction V (BSA), 2.0 g dextrose, 0.003 g catalase in 100 mL distilled water) at 37°C in an orbital shaker (100 rpm) until the OD600 reached up to 0.35. All procedures were performed at a BSL-3 facility. For further analysis, we used commercially available BSA Cohn fraction V (Merck, Darmstadt, Germany) unless otherwise stated. In addition, we used molecular biology grade Tween 80 (Merck product number P5188-100ML) without predilution.

### Counting the number of culturable cells

The cultured bacterial suspension was serially diluted 10-fold in phosphate buffered saline (pH 6.8) with 0.1% (w/v) Tween 80 (PBS-T) and 25 µL of each diluent was inoculated onto a Middlebrook 7H10 agar plate supplemented with oleic-albumin-dextrose-catalase (OADC) (Becton Dickinson, Sparks, MD) in duplication. Colonies were counted after at least three weeks of incubation at 37 °C under 5% CO2. The limit of detection was determined to be 20 CFU/mL owing to the diluent factor.

### Effect of BSA on the growth of Mtb cells in Dubos medium

Ten milliliters of log phase culture of Mtb H37Rv (OD600 = 0.35) in Dubos medium, as described previously, was sedimented at 3,000 ×*g* for 10 min at 4 °C, and the supernatant was discarded. The sediment was washed twice with fresh Dubos broth and resuspended in 10 mL Dubos broth. The suspension was diluted 1:10 with Dubos broth with or without the addition of BSA solution (5.0 g BSA fraction V in 100 mL distilled water) at a final concentration of 0.1% (w/v) BSA. The diluted suspension was dispensed into 24-well plates (Sarstedt, Nümbrecht, Germany) in 1.5 mL increments and sealed with gas-permeable film (Breathe-Easy, Diversified Biotech Inc, Dedham, MA, United States). Cultures were incubated for 5, 10, 15, and 20 days at 37 °C under 5% CO2. At each time point, the number of culturable cells was measured, as described above.

### Induction of the VBNC cells

Induction of the VBNC cells was performed according to a previous report (16) with some modifications. In short, the log phase culture of Mtb H37Rv or H37Ra (OD600 = 0.35) in Dubos medium was evenly divided into 30 mL PETG bottles, and 5 mg/mL diphenyleneiodonium chloride (DPI) in dimethyl sulfoxide (DMSO) solution was directly added at a final concentration of 4 µg/mL with immediate agitation. An equal volume of DMSO was added to the untreated control samples. The cultures were incubated for 24 h at 37 °C without shaking. After incubation, their culturability and viability were measured as described above.

### Cell viability assay

Mtb cells was measured by esterase activity and membrane integrity of the cells using dual staining with 10 mg/mL carboxyfluorescein diacetate (CFDA, DOJINDO LABORATORIES, Kumamoto, Japan) and ethidium bromide (EtBr, DOJINDO LABORATORIES). Briefly, 500 µL of DPI-treated or untreated Mtb culture was sedimented by centrifugation at 15,000 ×*g* for 5 min at 4 °C and washed twice with an equal volume of PBS-T. The washed bacterial sediment was resuspended in 200 µL of PBS-T with CFDA and EtBr and incubated at room temperature (RT) for 30 min in the shade. The final concentrations of CFDA and EtBr were 60 µg/mL and 2 µg/mL, respectively. Stained Mtb cells were washed twice with PBS-T and fixed with 3.7% formaldehyde solution at RT for at least 2 h. After fixation, 10 µL of the bacterial suspension was smeared on three APS-coated 8-well slides (Matsunami Glass, Osaka, Japan) and air-dried. The slides were then mounted with DPX new non-aqueous mounting medium (Merck, Darmstadt, Germany) and a 150 µm thick cover glass (Matsunami glass, Osaka, Japan) prior to microscopic observation. Specimens were examined using a BX53 fluorescence microscope (Olympus, Tokyo, Japan) with a 100× objective lens for green (esterase-active with intact membrane; live) or red (esterase-negative with damaged membrane; dead) fluorescing cells. At least 10 random fields were observed for each sample and the images (TIFF format) were analyzed using the Fiji/ImageJ software program(47). The live/dead ratio was calculated by directly counting green or red fluorescent cells.

### Auramine-O/Nile Red dual staining

Loss of acid fastness and accumulation of neutral lipids are distinctive features of VBNC mycobacteria(48). To detect the phenotype, fluorescent acid-fast staining with auramine-O and neutral lipid staining with Nile Red (9-dimethylamino-5H-benzo-α-phenoxadine-5-one) were performed using a previously described method (9) with some modifications. Ten microliters of DPI-treated or untreated cultures were smeared onto an APS-coated slide (Matsunami Glass, Osaka, Japan) and air-dried. The slides were then heat-fixed and cooled to room temperature before staining. The smear was flooded with fluorochrome staining solution (10 µg/mL auramine-O in 5% [w/v] phenol solution) and incubated at room temperature for 20 min in the dark. The excess dye was removed using 3% hydrochloric acid in ethanol for 15 min. The smear was then covered with 10 µg/mL Nile Red in ethanol and incubated at room temperature for 15 min. Finally, the smear was counterstained with a 0.1% (w/v) potassium permanganate solution for 1 min. The stained slides were air-dried and mounted, as previously described. Fluorescence microscopy was performed with a 100× objective lens for green (acid-fastness positive, lipid accumulation negative; active) or red (acid-fastness negative, lipid accumulation positive; VBNC) fluorescent cells. At least five random fields were observed for each sample, and images (TIFF format) were analyzed using the Fiji/ImageJ software program.(47) The proportion of acid-fast-positive cells and lipid-accumulation-positive cells was calculated by direct counting of green or red fluorescent cells.

### Acid-fast staining

Ziehl-Neelsen staining was also performed to confirm the loss of acid-fastness resulting in the VBNC state by DPI treatment, according to the standard method(49). The slides were observed under a light microscope with a 100× objective lens (BX53, Olympus, Tokyo, Japan).

### Confocal super-resolution microscopy

Detailed images of accumulated lipids were obtained using a confocal laser-scanning microscope (LSM900 with Airyscan 2, Carl Zeiss Microscopy, Jena, Germany). Briefly, culture slides of DPI-treated cells dual-stained with auramine-O and Nile Red, as described above, were observed with a 63× objective lens. The green fluorescence emission from auramine-O and red fluorescence from Nile Red were sequentially collected using a variable dichroic beam splitter and an Airyscan 2 detector. Super-resolution images were processed using ZEN Blue software (Carl Zeiss, Jena, Germany).

### VBNC reactivation assay

The VBNC reactivation assay was performed as described previously (16) with some modifications. DPI-treated cultures were sedimented by centrifugation at 3,000 x*g* for 10 min at 4 °C, washed twice with fresh Dubos broth, and resuspended in ten-milliliters of Dubos broth. The bacterial suspension was diluted 1:10 with fresh Dubos medium, with or without 0.1% (w/v) BSA. For both the BSA-containing and BSA-free series, OADC supplementation (Becton Dickinson, Sparks, MD, USA), fetal bovine serum (FBS; Moregate BioTech, Bulimba, Queensland, Australia), or sodium pyruvate (PA; Merck) were added. The final concentration of each reagent was 10% (v/v) for OADC supplementation, 2% (v/v) for FBS, and 3 mM for PA. The diluted suspension was then dispensed into 24-well plates in 1.5 mL increments and sealed with a gas-permeable film. Cultures were incubated for 5, 10, 15, and 20 days at 37 °C under 5% CO2. At each time point, the number of culturable cells was measured, as previously described.

### Evaluation of the reactivation-promoting effect of human and mouse serum albumin, egg-white albumin (ovalbumin) and antioxidative agents

To determine whether the reactivation-promoting effect of BSA is its antioxidative property, we measured the reactivation-promoting activity of other albumins that have variable antioxidative properties and antioxidative agents toward DPI-treated Mtb cells. Washed DPI-treated cells were resuspended in fresh Dubos broth without BSA, and the suspension was diluted 1:10 with Dubos broth containing BSA, human serum albumin (HSA), mouse serum albumin (MSA), ovalbumin (OVA), N-acetyl-*L-*cysteine (NAC), or D-mannitol (MAN). The final concentration of each reagent was 0.1% (w/v) for BSA, HSA, MSA, and OVA; 500 µM for NAC; and 50 mM for MAN. The subsequent reactivation assay was performed according to the method described above.

### Evaluation of the reactivation-promoting effect of fatty acid and globulin-free BSA

To determine the effect of BSA Cohn fraction V contaminants, we measured the reactivation-promoting activity of fatty acid and globulin-free BSA toward DPI-treated VBNC Mtb cells. Washed DPI-treated cells were resuspended in fresh Dubos broth without BSA, and the suspension was diluted 1:10 with Dubos broth containing BSA Cohn fraction V or fatty acid and globulin-free BSA. Fatty acid and globulin-free BSA were acquired from Merck. The final concentration of both the BSAs was 0.1% (w/v). The subsequent reactivation assay was performed according to the method described above.

### Effects of adenylyl cyclase or protein kinase on BSA-induced reactivation

Washed DPI-treated VBNC Mtb cells were resuspended in fresh Dubos broth, and the suspension was diluted to 1:10 with Dubos broth containing 0.1% (w/v) BSA containing adenylyl cyclase inhibitor SQ22536 (Merck, Darmstadt, Germany), protein kinase inhibitor H89 (Abcam, Cambridge, United Kingdom), or staurosporine (Merck). The final concentrations of each inhibitor were as follows: 0.1 to 10 mM SQ22536, 1–30 µM H89 and 1 to10 µM for staurosporine. The number of culturable cells was then measured as described previously.

### Molecular docking simulation

Molecular docking simulation was performed using the AutoDock Vina software program (version 1.1.2), an open-source molecular docking program designed by the Scripps Research Institute(50, 51). The 3D structures of the receptor proteins PknA and Rv*2212* were obtained from the Protein Data Bank (52) and AlphaFold Database(53, 54), respectively. The PDB ID for PknA was 4OW8 (55) and the AlphaFold Database identifier for Rv*2212* was AF-P9WMU7-F1. We removed water molecules from the proteins, added polar hydrogen and charge, and adjusted the X-Y-Z coordinates and grid size to optimize molecular docking. Each simulation was performed using default options. The 3D structures of the ligand molecules, H89, SQ22536 and ATP were obtained from the PubChem Database. The PubChem CIDs of the ligands were 449241, 5270, and 5957, respectively. Graphical processing was performed using the UCSF Chimera software program version 1.16(56).

### Statistical analysis

The statistical significance of differences between culturable cell counts of Mtb cells in untreated and DPI-treated cells was determined by Student’s *t*-test. The statistical significance of differences between staining of Mtb cells in untreated and DPI-treated cells was determined using Welch’s *t-*test. The statistical significance of differences between culturable cell counts of DPI-treated Mtb cells at days 0 and 20 on reactivation was determined by the Student’s *t-* test. For multiple group comparisons, one-way analysis of variance (ANOVA) was performed, followed by Tukey’s honest significant difference (HSD) test for multiple comparisons. Only comparisons with the control group (day 0) were extracted. Statistical analysis was performed with R (57) using EZR in the R package(58). *P* < 0.05.

## References

1. World Health Organization. 2020. TB disease burden, p. 23. In Global Tuberculosis Report 2020 Chapter 4. World Health Organization.

2. Ayrapetyan M, Williams T, Oliver JD. 2018. Relationship between the Viable but Nonculturable State and Antibiotic Persister Cells. Journal of Bacteriology 200.

3. Li L, Mendis N, Trigui H, Oliver JD, Faucher SP. 2014. The importance of the viable but non-culturable state in human bacterial pathogens. Frontiers in Microbiology 10.3389/fmicb.2014.00258.

4. Oliver JD. 2010. Recent findings on the viable but nonculturable state in pathogenic bacteria. FEMS Microbiology Reviews 34:415–425.

5. Wayne LG, Hayes LG. 1996. An in vitro model for sequential study of shiftdown of *Mycobacterium tuberculosis* through two stages of nonreplicating persistence. Infection and immunity 64:2062–9.

6. Betts JC, Lukey PT, Robb LC, McAdam RA, Duncan K. 2002. Evaluation of a nutrient starvation model of *Mycobacterium tuberculosis* persistence by gene and protein expression profiling. Molecular microbiology 43:717–31.

7. Salina EG, Waddell SJ, Hoffmann N, Rosenkrands I, Butcher PD, Kaprelyants AS. 2014. Potassium availability triggers *Mycobacterium tuberculosis* transition to, and resuscitation from, non-culturable (dormant) states. Open Biology 4:140106.

8. Aguilar-Ayala DA, Cnockaert M, Vandamme P, Palomino JC, Martin A, Gonzalez-Y-Merchand J. 2018. Antimicrobial activity against *Mycobacterium tuberculosis* under in vitro lipid-rich dormancy conditions. Journal of medical microbiology 67:282–285.

9. Deb C, Lee CM, Dubey VS, Daniel J, Abomoelak B, Sirakova TD, Pawar S, Rogers L, Kolattukudy PE. 2009. A novel in vitro multiple-stress dormancy model for *Mycobacterium tuberculosis* generates a lipid-loaded, drug-tolerant, dormant pathogen. PLoS ONE 4:e6077.

10. Desnues B, Cuny C, Grégori G, Dukan S, Aguilaniu H, Nyström T. 2003. Differential oxidative damage and expression of stress defence regulons in culturable and non-culturable *Escherichia coli* cells. EMBO reports 4:400–404.

11. Imazaki I, Nakaho K. 2009. Temperature-upshift-mediated revival from the sodium-pyruvate-recoverable viable but nonculturable state induced by low temperature in *Ralstonia solanacearum*: linear regression analysis. Journal of General Plant Pathology 2009 75:3 75:213–226.

12. Mizunoe Y, Wai SN, Ishikawa T, Takade A, Yoshida S. 2000. Resuscitation of viable but nonculturable cells of *Vibrio parahaemolyticus* induced at low temperature under starvation. FEMS Microbiology Letters 186:115–120.

13. Lynn M, Wilson AR, Solotorovsky M. 1979. Role of bovine serum albumin in the nutrition of *Mycobacterium tuberculosis*. Applied and Environmental Microbiology 38:806–810.

14. Dubos RJ. 1947. THE EFFECT OF LIPIDS AND SERUM ALBUMIN ON BACTERIAL GROWTH. The Journal of experimental medicine 85:9–22.

15. Davis BD, Dubos RJ. 1947. THE BINDING OF FATTY ACIDS BY SERUM ALBUMIN, A PROTECTIVE GROWTH FACTOR IN BACTERIOLOGICAL MEDIA. The Journal of experimental medicine 86:215–228.

16. Yeware A, Gample S, Agrawal S, Sarkar D. 2019. Using diphenyleneiodonium to induce a viable but non-culturable phenotype in *Mycobacterium tuberculosis* and its metabolomics analysis. PLoS ONE 14:e0220628.

17. Roche M, Rondeau P, Singh NR, Tarnus E, Bourdon E. 2008. The antioxidant properties of serum albumin. FEBS Letters 582:1783–1787.

18. Shleeva M, Goncharenko A, Kudykina Y, Young D, Young M, Kaprelyants A. 2013. Cyclic amp-dependent resuscitation of dormant mycobacteria by exogenous free fatty acids. PLoS ONE 8:e82914.

19. Shleeva MO, Kondratieva TK, Demina GR, Rubakova EI, Goncharenko A V., Apt AS, Kaprelyants AS. 2017. OVerexpression Of adenylyl cyclase encoded by the *Mycobacterium tuberculosis* Rv2212 gene confers improved fitness, accelerated recovery from dormancy and enhanced virulence in mice. Frontiers in Cellular and Infection Microbiology 7:370.

20. Knapp GS, Lyubetskaya A, Peterson MW, Gomes ALC, Ma Z, Galagan JE, McDonough KA. 2015. Role of intragenic binding of cAMP responsive protein (CRP) in regulation of the succinate dehydrogenase genes Rv0249c-Rv0247c in TB complex mycobacteria. Nucleic acids research 43:5377–93.

21. Morishige Y, Fujimori K, Amano F. 2013. Differential resuscitative effect of pyruvate and its analogues on VBNC (Viable But Non-Culturable) *Salmonella*. Microbes and Environments 28:180– 186.

22. Vilhena C, Kaganovitch E, Grünberger A, Motz M, Forné I, Kohlheyer D, Jung K. 2019. Importance of Pyruvate Sensing and Transport for the Resuscitation of Viable but Nonculturable *Escherichia coli* K-12. Journal of bacteriology 201.

23. Yoon JH, Bae YM, Jo S, Moon SK, Oh SW, Lee SY. 2020. Optimization of resuscitation-promoting broths for the revival of *Vibrio parahaemolyticus* from a viable but nonculturable state. Food Science and Biotechnology 30:159.

24. Ortega C, Liao R, Anderson LN, Rustad T, Ollodart AR, Wright AT, Sherman DR, Grundner C. 2014. *Mycobacterium tuberculosis* Ser/Thr Protein Kinase B Mediates an Oxygen-Dependent Replication Switch. PLOS Biology 12:e1001746.

25. Lyon RH, Lichstein HC, Hall WH. 1963. EFFECT OF TWEEN 80 ON THE GROWTH OF TUBERCLE BACILLI IN AERATED CULTURES. Journal of Bacteriology 86:280–284.

26. Huang BX, Kim HY, Dass C. 2004. Probing three-dimensional structure of bovine serum albumin by chemical cross-linking and mass spectrometry. Journal of the American Society for Mass Spectrometry 15:1237–1247.

27. Rombouts I, Lagrain B, Scherf KA, Lambrecht MA, Koehler P, Delcour JA. 2015. Formation and reshuffling of disulfide bonds in bovine serum albumin demonstrated using tandem mass spectrometry with collision-induced and electron-transfer dissociation. Scientific reports 5:12210.

28. Broersen K, Van Teeffelen AMM, Vries A, Voragen AGJ, Hamer RJ, De Jongh HHJ. 2006. Do sulfhydryl groups affect aggregation and gelation properties of ovalbumin? Journal of agricultural and food chemistry 54:5166–74.

29. Wells AQ. 1937. TUBERCULOSIS IN WILD VOLES. The Lancet 229:1221.

30. Manuse S, Fleurie A, Zucchini L, Lesterlin C, Grangeasse C. 2016. Role of eukaryotic-like serine/threonine kinases in bacterial cell division and morphogenesis. FEMS microbiology reviews 40:41–56.

31. Carette X, Platig J, Young DC, Helmel M, Young AT, Wang Z, Potluri L-P, Moody CS, Zeng J, Prisic S, Paulson JN, Muntel J, Madduri AVR, Velarde J, Mayfield JA, Locher C, Wang T, Quackenbush J, Rhee KY, Moody DB, Steen H, Husson RN. 2018. Multisystem Analysis of *Mycobacterium tuberculosis* Reveals Kinase-Dependent Remodeling of the Pathogen-Environment Interface. mBio 9.

32. Thakur M, Chakraborti PK. 2008. Ability of PknA, a mycobacterial eukaryotic-type serine/threonine kinase, to transphosphorylate MurD, a ligase involved in the process of peptidoglycan biosynthesis. The Biochemical journal 415:27–33.

33. Parikh A, Verma SK, Khan S, Prakash B, Nandicoori VK. 2009. PknB-mediated phosphorylation of a novel substrate, N-acetylglucosamine-1-phosphate uridyltransferase, modulates its acetyltransferase activity. Journal of molecular biology 386:451–64.

34. Jagtap PKA, Soni V, Vithani N, Jhingan GD, Bais VS, Nandicoori VK, Prakash B. 2012. Substrate-bound crystal structures reveal features unique to *Mycobacterium tuberculosis* N-acetyl-glucosamine 1-phosphate uridyltransferase and a catalytic mechanism for acetyl transfer. The Journal of biological chemistry 287:39524–37.

35. Gee CL, Papavinasasundaram KG, Blair SR, Baer CE, Falick AM, King DS, Griffin JE, Venghatakrishnan H, Zukauskas A, Wei J-R, Dhiman RK, Crick DC, Rubin EJ, Sassetti CM, Alber T. 2012. A phosphorylated pseudokinase complex controls cell wall synthesis in mycobacteria. Science signaling 5:ra7.

36. Kang C-M, Nyayapathy S, Lee J-Y, Suh J-W, Husson RN. 2008. Wag31, a homologue of the cell division protein DivIVA, regulates growth, morphology and polar cell wall synthesis in mycobacteria. Microbiology (Reading, England) 154:725–735.

37. Gupta M, Sajid A, Sharma K, Ghosh S, Arora G, Singh R, Nagaraja V, Tandon V, Singh Y. 2014. HupB, a nucleoid-associated protein of *Mycobacterium tuberculosis*, is modified by serine/threonine protein kinases in vivo. Journal of bacteriology 196:2646–57.

38. Baronian G, Ginda K, Berry L, Cohen-Gonsaud M, Zakrzewska-Czerwińska J, Jakimowicz D, Molle V. 2015. Phosphorylation of *Mycobacterium tuberculosis* ParB participates in regulating the ParABS chromosome segregation system. PloS one 10:e0119907.

39. Thakur M, Chakraborti PK. 2006. GTPase activity of mycobacterial FtsZ is impaired due to its transphosphorylation by the eukaryotic-type Ser/Thr kinase, PknA. The Journal of biological chemistry 281:40107–13.

40. Sureka K, Hossain T, Mukherjee P, Chatterjee P, Datta P, Kundu M, Basu J. 2010. Novel role of phosphorylation-dependent interaction between FtsZ and FipA in mycobacterial cell division. PloS one 5:e8590.

41. Wright DP, Ulijasz AT. 2014. Regulation of transcription by eukaryotic-like serine-threonine kinases and phosphatases in gram-positive bacterial pathogens. Virulence 5:863–885.

42. Rock JM, Hopkins FF, Chavez A, Diallo M, Chase MR, Gerrick ER, Pritchard JR, Church GM, Rubin EJ, Sassetti CM, Schnappinger D, Fortune SM. 2017. Programmable transcriptional repression in mycobacteria using an orthogonal CRISPR interference platform. Nature microbiology 2:16274.

43. Minato Y, Gohl DM, Thiede JM, Chacón JM, Harcombe WR, Maruyama F, Baughn AD. 2019. Genomewide Assessment of *Mycobacterium tuberculosis* Conditionally Essential Metabolic Pathways. mSystems 4.

44. DeJesus MA, Gerrick ER, Xu W, Park SW, Long JE, Boutte CC, Rubin EJ, Schnappinger D, Ehrt S, Fortune SM, Sassetti CM, Ioerger TR. 2017. Comprehensive Essentiality Analysis of the *Mycobacterium tuberculosis* Genome via Saturating Transposon Mutagenesis. mBio 8.

45. Sassetti CM, Boyd DH, Rubin EJ. 2003. Genes required for mycobacterial growth defined by high density mutagenesis. Molecular microbiology 48:77–84.

46. Griffin JE, Gawronski JD, Dejesus MA, Ioerger TR, Akerley BJ, Sassetti CM. 2011. High-resolution phenotypic profiling defines genes essential for mycobacterial growth and cholesterol catabolism. PLoS pathogens 7:e1002251.

47. Schindelin J, Arganda-Carreras I, Frise E, Kaynig V, Longair M, Pietzsch T, Preibisch S, Rueden C, Saalfeld S, Schmid B, Tinevez J-Y, White DJ, Hartenstein V, Eliceiri K, Tomancak P, Cardona A. 2012. Fiji: an open-source platform for biological-image analysis. Nature Methods 2012 9:7 9:676–682.

48. Garton NJ, Christensen H, Minnikin DE, Adegbola RA, Barer MR. 2002. Intracellular lipophilic inclusions of mycobacteria in vitro and in sputum. Microbiology 148:2951–2958.

49. Weyer K. 1998. Laboratory services in tuberculosis control. Part II. WHO Tech Bull 98:258.

50. Trott O, Olson AJ. 2010. AutoDock Vina: improving the speed and accuracy of docking with a new scoring function, efficient optimization, and multithreading. Journal of computational chemistry 31:455–61.

51. Eberhardt J, Santos-Martins D, Tillack AF, Forli S. 2021. AutoDock Vina 1.2.0: New Docking Methods, Expanded Force Field, and Python Bindings. Journal of chemical information and modeling 61:3891–3898.

52. Berman HM, Westbrook J, Feng Z, Gilliland G, Bhat TN, Weissig H, Shindyalov IN, Bourne PE. 2000. The Protein Data Bank. Nucleic Acids Research 28:235–242.

53. Jumper J, Evans R, Pritzel A, Green T, Figurnov M, Ronneberger O, Tunyasuvunakool K, Bates R, Žídek A, Potapenko A, Bridgland A, Meyer C, Kohl SAA, Ballard AJ, Cowie A, Romera-Paredes B, Nikolov S, Jain R, Adler J, Back T, Petersen S, Reiman D, Clancy E, Zielinski M, Steinegger M, Pacholska M, Berghammer T, Bodenstein S, Silver D, Vinyals O, Senior AW, Kavukcuoglu K, Kohli P, Hassabis D. 2021. Highly accurate protein structure prediction with AlphaFold. Nature 2021 596:7873 596:583–589.

54. Varadi M, Anyango S, Deshpande M, Nair S, Natassia C, Yordanova G, Yuan D, Stroe O, Wood G, Laydon A, Zídek A, Green T, Tunyasuvunakool K, Petersen S, Jumper J, Clancy E, Green R, Vora A, Lutfi M, Figurnov M, Cowie A, Hobbs N, Kohli P, Kleywegt G, Birney E, Hassabis D, Velankar S. 2022. AlphaFold Protein Structure Database: massively expanding the structural coverage of protein-sequence space with high-accuracy models. Nucleic Acids Research 50:D439–D444.

55. Ravala SK, Singh S, Yadav GS, Kumar S, Karthikeyan S, Chakraborti PK. 2015. Evidence that phosphorylation of threonine in the GT motif triggers activation of PknA, a eukaryotic-type serine/threonine kinase from *Mycobacterium tuberculosis*. The FEBS journal 282:1419–31.

56. Pettersen EF, Goddard TD, Huang CC, Couch GS, Greenblatt DM, Meng EC, Ferrin TE. 2004. UCSF Chimera - A visualization system for exploratory research and analysis. Journal of Computational Chemistry 25:1605–1612.

57. R Core Team. 2021. R: A Language and Environment for Statistical Computing. 4.1.0. R Foundation for Statistical Computing, Vienna, Austria.

58. Kanda Y. 2013. Investigation of the freely available easy-to-use software “EZR” for medical statistics. Bone marrow transplantation 48:452–8.

