## Supplementary material for "Human and bovine serum albumin, but not mouse serum and egg-white albumin, promote reactivation of viable but non-culturable *Mycobacterium tuberculosis* via the activation of protein kinase-dependent cell division processes": suppl_revised

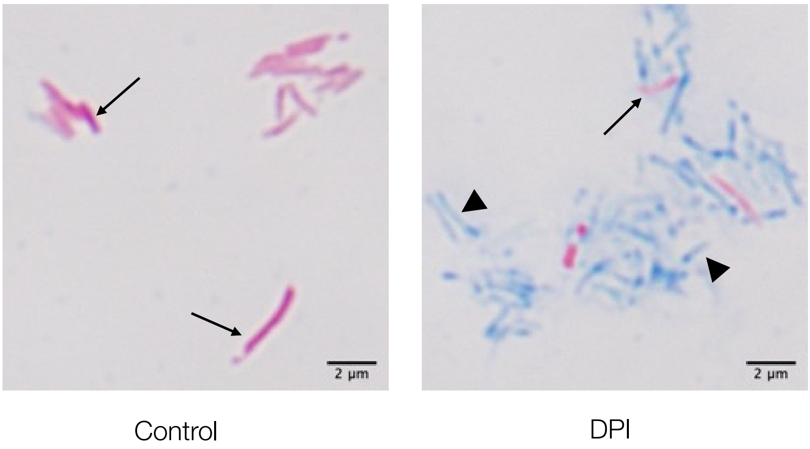


**Supplementary Figure 1**. Ziehl-Neelsen staining of DPI-treated Mtb
Slides were prepared according to the standard method. Red cells stained with carbol fuchsin (arrow) are acid-fast-positive, and blue cells stained with methylene blue (arrowhead) are acid-fast-negative.

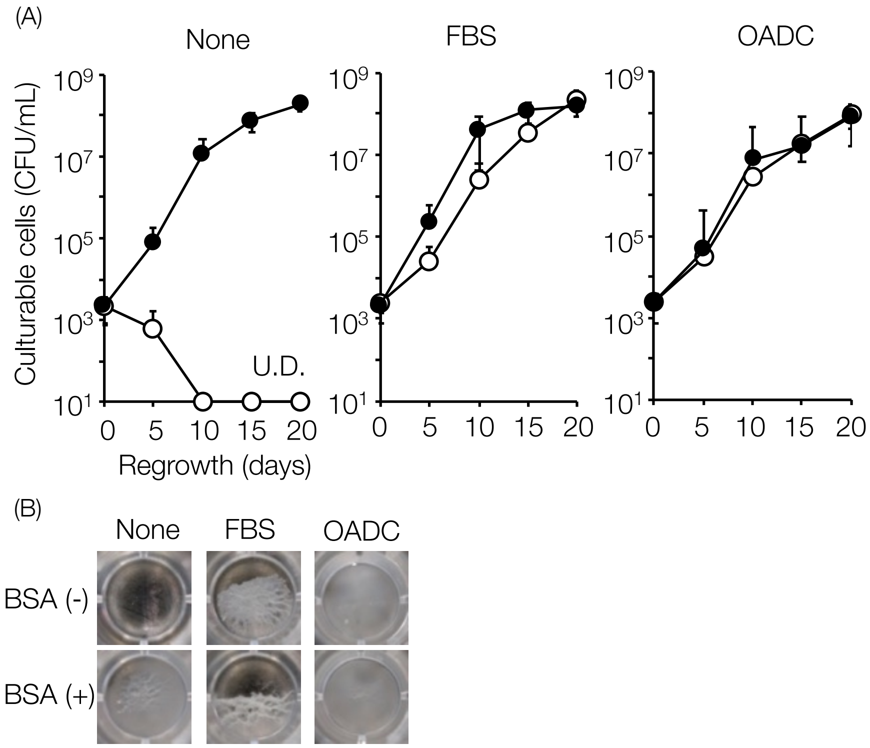


**Supplementary Figure 2.** Reactivation of DPI-treated Mtb H37Ra bacilli by FBS and OADC supplementation

(A) The number of culturable Mtb cells without supplementation or those cells supplemented with 2% FBS or 10% OADC in the presence (closed circle) or absence (open circle) of 0.1% BSA in DPI-treated cells at the indicated days of incubation. The CFU/mL values were determined by plating the cells onto 7H10 plates in duplicate. Data represent mean ± SD from three independent experiments. The U.D. was below the limit of detection (20 CFU/mL).

(B) Representative images of regrowth under the indicated conditions were captured at the end of incubation. None, FBS, and OADC represent no supplementation, 2% FBS supplementation, and 10% OADC supplementation, respectively.

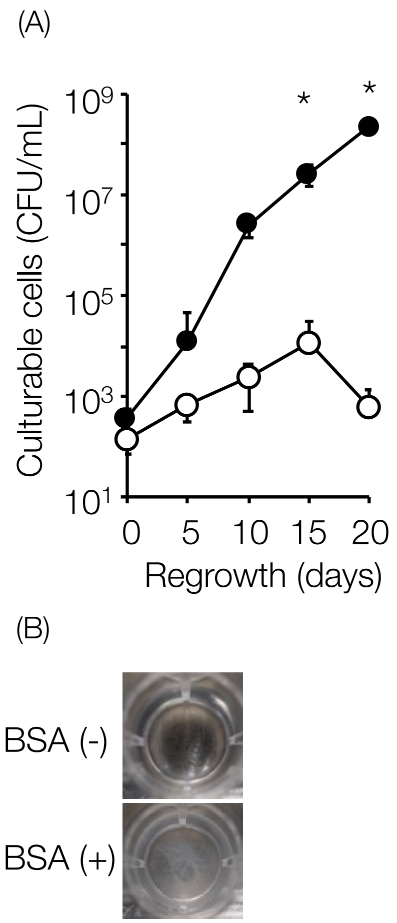


**Supplementary Figure 3.** Reactivation of DPI-treated Mtb H37Rv by pyruvate supplementation

(A) The number of culturable Mtb cells supplemented with 3 mM sodium pyruvate in the presence (closed circle) or absence (open circle) of BSA in DPI-treated cells at the indicated days of incubation. The CFU/mL values were determined by plating the cells onto 7H10 plates in duplicate. Data represent mean ± SD from three independent experiments. Asterisks indicate a statistically significant difference between the presence and absence of 0.1% BSA using Student's *t*-test. (*$p$<0.05)

(B) Representative images of regrowth under the indicated conditions were captured at the end of incubation.

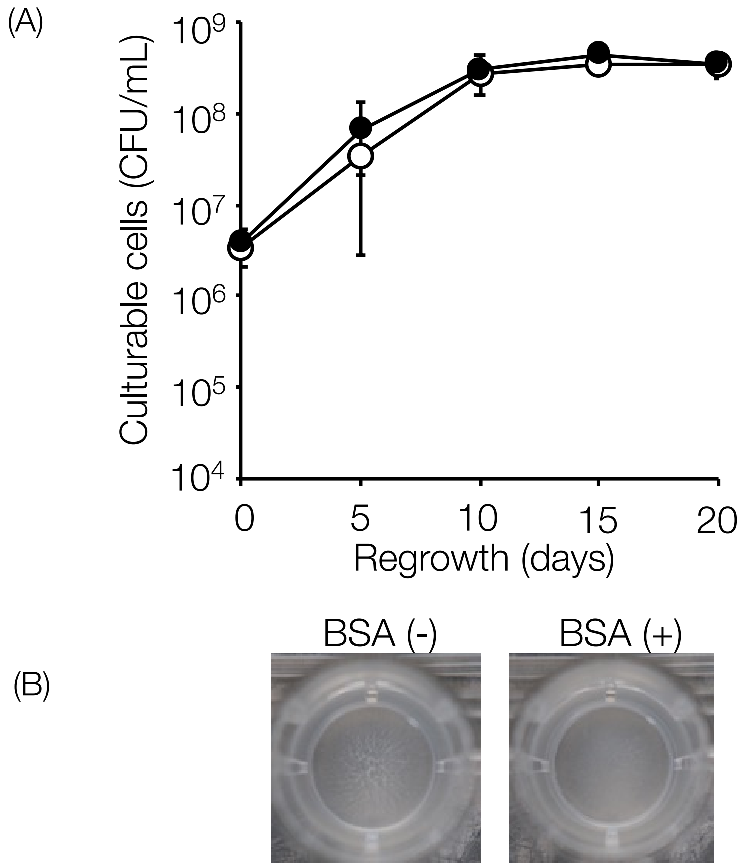


**Supplementary Figure 4.** Growth curves of Mtb H37Rv cells in Dubos' medium in the presence or absence of BSA

(A) Number of culturable cells in Dubos medium in the presence (closed circle) or absence (open circle) of 0.1% BSA on the indicated days of incubation. The CFU/mL values were determined by plating the cells onto 7H10 plates in duplicate. Data represent the mean ± SD from three independent experiments.

(B) Representative images of Mtb growth were captured at the end of the incubation (day 20).

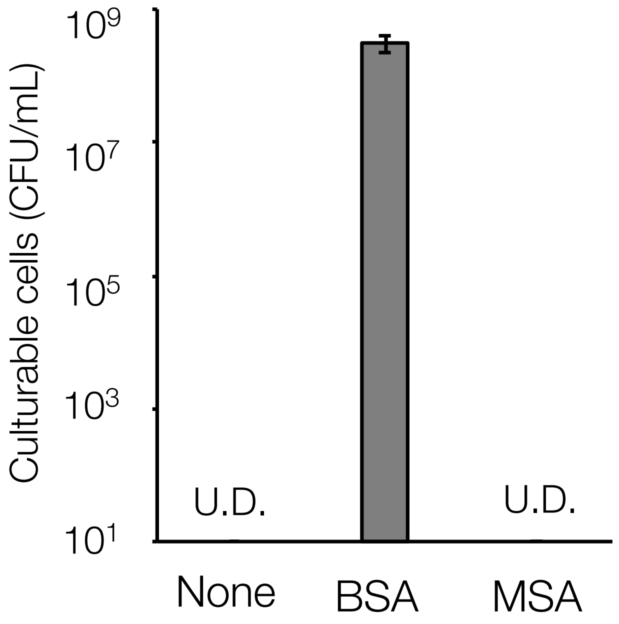


**Supplementary Figure 5.** Differential reactivation-promoting effect of BSA and mouse serum albumin (MSA) on the regrowth of DPI-treated Mtb

Number of culturable DPI-treated Mtb cells supplemented with 0.1% (w/v) BSA and MSA at the end of incubation. The CFU/mL value was determined by plating the cells onto a 7H10 plate in duplicate. Data are presented as the mean ± SD from three independent experiments.

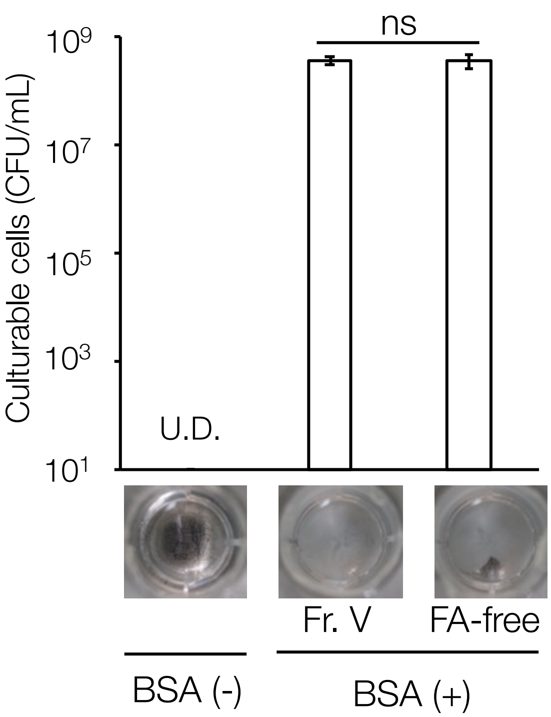


**Supplementary Figure 6.** Effect of fatty acid-free BSA on regrowth of DPI-treated Mtb

The number of culturable DPI-treated Mtb cells supplemented with 0.1% (w/v) bovine serum albumin (BSA) Cohn fraction V (Fr. V), 0.1% (w/v) fatty acid, and globulin-free BSA (FA-free) at the end of the incubation. CFU/mL values were determined by plating the cells onto 7H10 plates in duplicate. Data represent mean ± SD from three independent experiments. Representative images of Mtb regrowth captured at the end of incubation are shown in the figure.

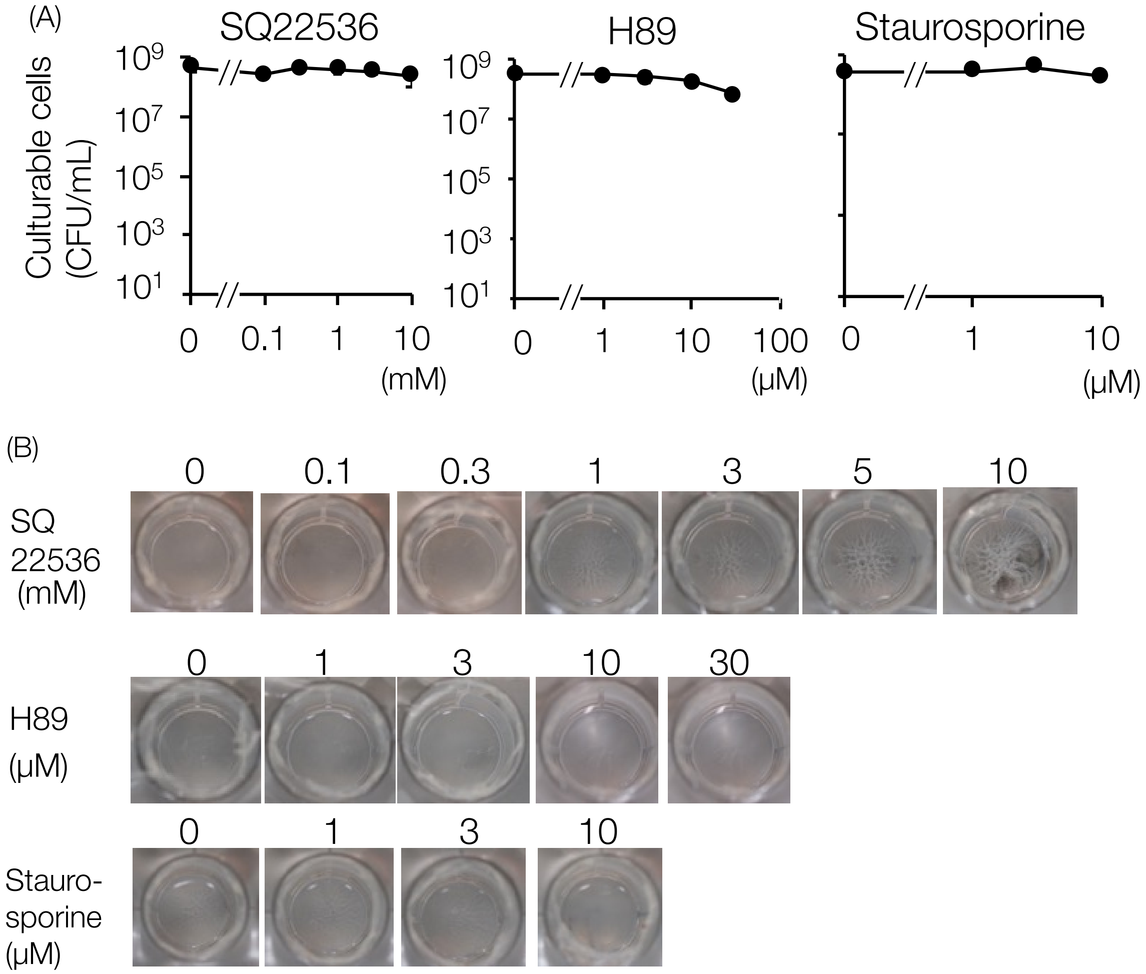


**Supplementary Figure 7.** Effect of SQ22536, H89, and staurosporine on the growth of intact, non-DPI-treated Mtb

(A) The culturable Mtb cells at the end of incubation were supplemented with 0.1% BSA, SQ22536, H89, or staurosporine. Inhibitors were added at the indicated concentrations.

(B) Representative images of Mtb regrowth were captured at the end of incubation (day 20).

The CFU/mL values were determined by plating the cells onto a 7H10 plate in duplicate. Data represent mean ± SD from three independent experiments.

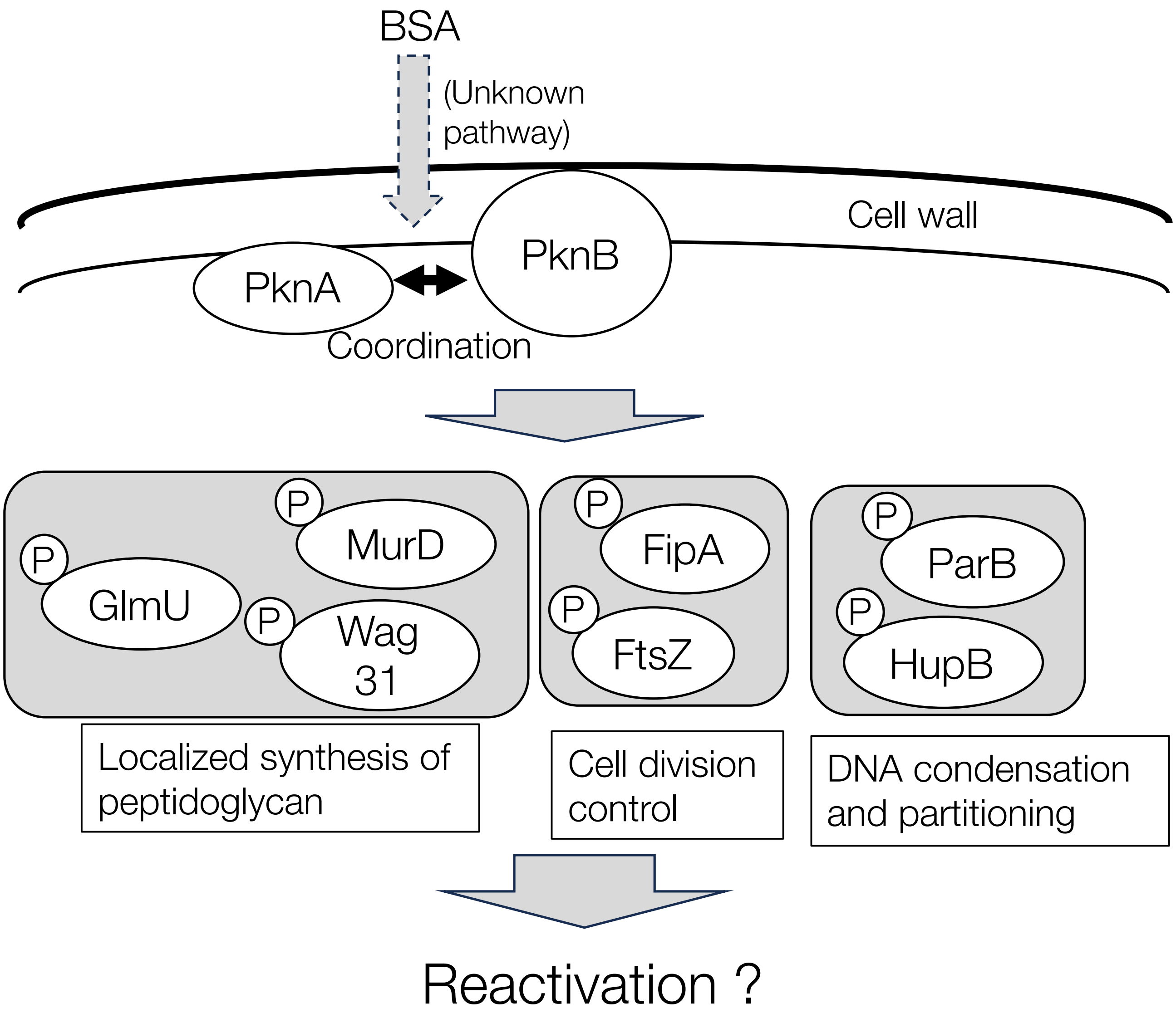


**Supplementary Figure 8.** Schematic representation of the mechanism of BSA-induced Mtb reactivation. Although the detailed mechanism of interaction between BSA and PknA/PknB is not yet clear, BSA may activate PknA/PknB and positively regulate target proteins involved in the localized synthesis of peptidoglycan, cell division control, and DNA condensation and partitioning through their phosphorylation, resulting in the reactivation of VBNC Mtb cells. The target proteins of PknA/PknB were predicted using the STRING Database (https://string-db.org/).

| Receptor | Ligand | Affinity (kcal/mol) | Best-docked complex |
|---|---|---|---|
| Rv2212 | SQ22536 | -5.5 | 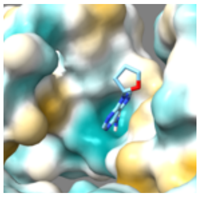 |
| | ATP | -6.9 | 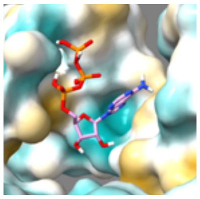 |
| PknA | H89 | -7.9 | 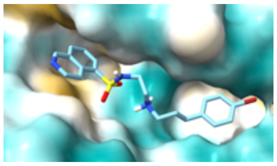 |
| | ATP | -7.5 | 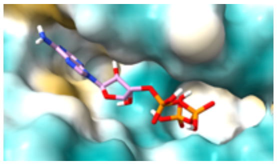 |

**Supplementary Table 1.** Molecular docking results

Affinity was calculated using Autodock Vina software. "Best-docked complex" shows images of the best docking of the receptor and ligand. The receptor surface was colored according to amino acid hydrophobicity: cyan for the most hydrophilic residues, white for neutral residues, and yellow for the most hydrophobic residues.
